# Multimodal Image Normalisation Tool (MINT) for the Adolescent Brain and Cognitive Development study: the MINT ABCD Atlas

**DOI:** 10.1101/2022.08.09.503395

**Authors:** Diliana Pecheva, John R Iversen, Clare E Palmer, Richard Watts, Terry L Jernigan, Donald J Hagler, Anders M Dale

## Abstract

The Adolescent Brain and Cognitive Development (ABCD) study aims to measure the trajectories of brain, cognitive, and emotional development. Cognitive and behavioural development during late childhood and adolescence have been associated with a myriad of microstructural and morphological alterations across the brain, as measured by magnetic resonance imaging (MRI). These associations may be strongly localised or spatially diffuse, therefore, it would be advantageous to analyse multimodal MRI data in concert, and across the whole brain. The ABCD study presents the unique challenge of integrating multimodal data from tens of thousands of scans at multiple timepoints, within a reasonable computation time. To address the need for a multimodal registration and atlas for the ABCD dataset, we present the synthesis of an ABCD atlas using the Multimodal Image Normalisation Tool (MINT). The MINT ABCD atlas was generated from baseline and two-year follow up imaging data using an iterative approach to synthesise a cohort-specific atlas from linear and nonlinear deformations of eleven channels of diffusion and structural MRI data. We evaluated the performance of MINT against two widely used methods and show that MINT achieves comparable alignment to current state-of-the-art multimodal registration, at a fraction of the computation time. To validate the use of the ABCD MINT atlas in whole brain, voxelwise analysis, we replicate and expand on previously published region-of-interest analysis between diffusion MRI-derived measures and body mass index (BMI). We also report novel association between BMI and brain morphology derived from the registration deformations. We present the ABCD MINT atlas as a publicly available resource to facilitate whole brain voxelwise analyses for the ABCD study.

## Introduction

Late childhood and adolescence are periods of substantial cognitive and behavioural development and are associated with the emergence of many psychiatric disorders. The Adolescent Brain and Cognitive Development (ABCD) study aims to measure the trajectories of brain, cognitive, and emotional development, among numerous other developmental outcomes, and to identify the genetic and environmental factors that may influence them. It is the largest long-term study of brain development and child health in the United States (Casey et al., 2018), and is epidemiologically-informed, including participants from demographically diverse backgrounds (Dick et al., 2021; Garavan et al., 2018).

Developmental and behavioural changes, both in typical and atypical development, have been associated with a myriad of microstructural and morphological alterations across the brain, as measured by magnetic resonance imaging (MRI) (Ball et al., 2018; Bos et al., 2018; Casey et al., 2019; Casey et al., 2005; Genc et al., 2017; Lenroot et al., 2007; Palmer et al., 2021). MRI modalities vary in their sensitivity to individual differences in behavioural development and in the trajectories of their derived measures (Bethlehem et al., 2022; Brown et al., 2012; Fjell et al., 2015; Lebel and Beaulieu, 2011; Lebel and Deoni, 2018). Furthermore, developmental brain-behaviour associations may be strongly localised, driven by particular anatomical features, or diffuse across the whole brain. It would therefore be advantageous for developmental studies to analyse multimodal MRI data in concert, and across the whole brain.

The first step in multimodal, whole brain, voxelwise analyses is to establish anatomical correspondence between participants, across all scans. This requires spatial normalisation of scans to a common space, such as an atlas. It is necessary to use a template that is an adequate representative of the overall cohort (Keihaninejad et al., 2012; Van Hecke et al., 2011; Yoon et al., 2009; Zhang and Arfanakis, 2013). Another consideration is the choice of MRI modality used to align scans. Diffusion MRI (dMRI) data contain rich orientational information which is best suited to aligning white matter and oriented structures. Structural MRI (sMRI) data offer higher resolution and greater grey matter-white matter contrast and are best suited to aligning cortical and subcortical structures. As different modalities can provide complementary information, the ideal registration algorithm would be able to leverage multiple channels of information. Most recently published paediatric atlases fall short of at least one of these stipulations and therefore are not ideal for the ABCD study. They either include too few participants, do not include demographically diverse participants, or were constructed using unimodal registration (Avants et al., 2015; Fonov et al., 2011; Molfese et al., 2021; Morris et al., 2020; Sanchez et al., 2012; Wilke et al., 2008; Xie et al., 2015; Yoon et al., 2009; Zhao et al., 2019; Zhu et al., 2021) (see Supplementary Table 1 for a summary).

The ABCD cohort includes 11,880 children being followed on a regular basis from age 9-10 years, for at least 10 years, with multimodal neuroimaging data collected every 2 years. This presents the unique challenge of integrating multimodal data from tens of thousands of scans at multiple timepoints, with consistently good alignment across participants, within a reasonable computation time. One widely used method that does employ a multimodal framework for registration and can be used to generate cohort-specific atlases is ANTs (Avants et al., 2008; Avants et al., 2014). While generally accepted to produce state-of-the-art alignment across participants and modalities (Klein et al., 2009), ANTs is computationally intensive. Computation time becomes prohibitively long when considering large datasets with tens of thousands of scans.

To address the need for a multimodal registration and atlas for the ABCD dataset, we present the synthesis of an ABCD atlas using the Multimodal Image Normalisation Tool (MINT). The MINT ABCD atlas is generated from baseline and two-year follow up imaging data. We compare the performance of MINT with that of two widely used methods, ANTs and FSL’s FLIRT/FNIRT. To validate the use of the ABCD MINT atlas in a whole brain, voxelwise analysis, we reproduced previously reported associations between dMRI-derived measures and body mass index (BMI) in the ABCD dataset.

The ABCD MINT atlas will be publicly available shortly from our GitHub repository at https://github.com/cmig-research-group/cmig_tools.

## Methods

### Sample

This paper uses baseline and two-year follow up (FU) data from the NIMH Data Archive ABCD Collection Release 4.0 (DOI: 10.15154/1523041). The ABCD cohort is epidemiologically informed (Garavan et al., 2018); it includes participants from demographically diverse backgrounds and has an embedded twin cohort and many siblings. Exclusion criteria for participation in the ABCD Study were limited to: 1) lack of English proficiency in the child; 2) the presence of severe sensory, neurological, medical or intellectual limitations that would inhibit the child’s ability to comply with the protocol; 3) an inability to complete an MRI scan at baseline. The study protocols are approved by the University of California, San Diego Institutional Review Board. Parent/caregiver permission and child assent were obtained from each participant. In total, 18,718 scans were inputted to the registration and atlas construction. The demographics of the sample are summarised in Table 1.

**Table 1.** Summary demographics of the sample of observations used in the atlas construction.

|  | Baseline | Two-year FU |
| --- | --- | --- |
| n | 11140 | 7578 |
| Age, median years (range) | 9.9 (8.9-11) | 11.9 (10.6-13.8) |
| Male sex, n (%) | 5787 (51.9%) | 4075 (54.8) |
| *Parental education* | n (%) | n (%) |
| < HS diploma, | 538 (4.8) | 316 (4.2) |
| HS diploma/GED | 1028 (9.2) | 664 (8.8) |
| Some college | 2870 (25.8) | 1881 *24.80 |
| Bachelor degree | 2847 (25.6) | 1988 (26.2) |
| Postgraduate degree | 3844 (34.5) | 2708 (35.7) |
| NA | 13 (0.1) | 86 (1.1) |
| *Race (race.6level)* | n (%) | n (%) |
| AIAN/NHPI | 75 (0.7) | 56 (0.7) |
| Asian | 253 (2.3) | 152 (2) |
| Black | 1698 (15.2) | 1039 (13.7) |
| Mixed | 1332 (12.0) | 914 (12.1) |
| Other | 485 (4.4) | 310 (4.1) |
| White | 7138 (64.1) | 5017 *66.2) |
| NA | 159 (1.4) | 90 (1.2) |
| Hispanic ethnicity | n (%) | n (%) |
| Yes | 2261 (20.3%) | 1460 (19.3) |
| No | 8733 (78.4) | 6032 (79.6) |
| NA | 146 (1.3) | 86 (1.1) |
| *Household income* | n (%) | n (%) |
| <50k | 2957 (26.5) | 1680 (22.2) |
| 50-100k | 2882 (25.9) | 1963 (25.9) |
| >100k | 4358 (39.1) | 3339 (44.1) |
| NA | 943 (8.5) | 596 (7.9) |

### Data Acquisition

T1-weighted (T1w) and dMRI data were collected using Siemens Prisma and Prisma Fit, General Electric MR Discovery 750, and Philips Achieva and Ingenia 3T scanners. Scanning protocols were harmonised across 21 sites. Full details of the image acquisition protocols used in the ABCD study have been described by Casey et al (2018) and Hagler et al (2019), therefore only a short overview is given here. dMRI data were acquired in the axial plane at 1.7 mm isotropic resolution with multiband acceleration factor 3 (except for the Ingenia, which used acceleration factor 4). Diffusion-weighted images were collected with seven b=0 s/mm^2^ frames and 96 non-collinear gradient directions, with 6 directions at b=500 s/mm^2^, 15 directions at b=1000 s/mm^2^, 15 directions at b=2000 s/mm^2^, and 60 directions at b=3000 s/mm^2^. T1w images were acquired using a 3D magnetisation-prepared rapid acquisition gradient echo (MPRAGE) scan with 1mm isotropic resolution and no multiband acceleration.

### Image preprocessing

#### dMRI

Imaging data were preprocessed using the ABCD Study processing pipeline as described by Hagler et al (2019). dMRI data were corrected for eddy current distortion using a diffusion gradient model-based approach (Zhuang et al., 2006). Head motion was corrected by rigid body registration of each diffusion-weighted volume to a corresponding volume synthesised from a robust tensor fit, accounting for image contrast variation between frames. Dark slices caused by abrupt head motion were replaced with values synthesised from the robust tensor fit, and the diffusion gradient matrix was adjusted for head rotation (Hagler et al., 2019). Spatial and intensity distortions caused by B0 field inhomogeneity were corrected using FSL’s *topup* (Andersson et al., 2003), followed by gradient nonlinearity distortion correction (Jovicich et al., 2006). The dMRI data were registered to T1w structural images using mutual information (Wells et al., 1996) after coarse pre-alignment via within-modality registration to atlas brains. dMRI data were then resampled to 1.7 mm isotropic resolution, equal to the dMRI acquisition resolution.

The restriction spectrum imaging (RSI) model (Brunsing et al., 2017; White et al., 2013a; White et al., 2014; White et al., 2013b) was fit to the diffusion-weighted images. In general, RSI is a flexible framework for modelling contributions of separable pools of water within tissues to the measured diffusion signal. As applied in ABCD, RSI is a three compartment model used to estimate the relative contribution of restricted, hindered and free water environments within brain tissue. A more detailed explanation of RSI and its derived measures as applied in ABCD is given by Palmer et al. (2021). Briefly, spherical deconvolution (SD) is used to reconstruct a fiber orientation distribution (FOD) in each voxel for each compartment. The hindered and restricted compartments are modelled as fourth order spherical harmonic coefficients (SH) functions, and the free water compartment is modelled using zeroth order SH functions. The axial diffusivity (AD) is held constant, with a value of 1×10^−3^ mm^2^/s for the restricted and hindered compartments. For the restricted compartment, the radial diffusivity (RD) is fixed to 0 mm^2^/s. For the hindered compartment, RD is fixed to 0.9×10^−3^ mm^2^/s. For the free water compartment the isotropic diffusivity is fixed to 3×10^−3^ mm^2^/s.

For each participant, major WM tracts were labelled using AtlasTrack, a probabilistic atlas-based method for automated segmentation of WM fiber tracts (Hagler et al., 2009; Hagler et al., 2019), and binarised.

#### sMRI

T1w images were preprocessed according to the standard ABCD processing pipeline, outlined in detail in Hagler et al (2019). T1w images were corrected for gradient nonlinearity distortions using scanner-specific, nonlinear transformations provided by MRI scanner manufacturers (Jovicich et al., 2006). Intensity inhomogeneity correction was performed by applying a smoothly varying, estimated B1-bias field (Hagler et al., 2019). Images were rigidly registered and resampled into alignment with a pre-existing, in-house, averaged, reference brain with 1.0 mm isotropic resolution (Hagler et al., 2019).

For each participant, cortical and subcortical grey matter structures were labelled using the Freesurfer 7.1.1 segmentation (Fischl et al., 2002).

### MINT registration

Images were registered according to the numerical method outlined by Holland et al. (2011), extended to include multimodal inputs. The registration algorithm consisted of rigid body, affine and nonlinear transformations. Eleven multimodal channels were used to align scans and create the MINT ABCD atlas, however the input channels varied according to the registration step. Three sMRI channels were included: T1w images, white matter segmentation, and grey matter segmentation. Eight dMRI channels were included: the zeroth and second order SH coefficients of the restricted FOD and the zeroth order SH coefficients from the hindered and free water FODs from the RSI model. Prior to the multimodal registration it is necessary to align the different modalities within each participant. This was achieved using the rigid body registration transforms from the ABCD processing pipeline (Hagler et al., 2019). A combined image of the white matter and grey matter segmentations was used as a single input channel for the two linear registration steps. This was done because rigid body and affine transformations and contain no local deformations, therefore only information regarding the size, orientation and positioning of the brain volume is relevant at these stages. Conversely, all eleven channels were input for the nonlinear registration step as this involves local deformations for which local differences in intensity are necessary to align anatomical structures.

The atlas-building algorithm was implemented as proposed by Joshi et al (2004), such that the registration target was the group mean image, constructed iteratively, and transformation fields were estimated for all participants towards the provisional group mean.

#### Linear registration

After preprocessing, moving images are registered to a target image using rigid body and then affine transformations. A moving image is rigid body transformed, via a transformation with 6 degrees of freedom (rotation and translation parameters), to align with the target image. Images are registered by minimising a cost function based on intensity differences between the moving and target images in voxels within a brain mask. The mask is defined in atlas space from the group average b=0 mm/s^2^ image. Images are masked such that the values of the mask are 1 inside the brain and fall off to 0 outside the brain along a spatially smooth gradient of several voxels to avoid artificially introducing sharp boundaries in the images. Alignment should be driven by brain structure and using this type of mask downweighs the effects of non-brain tissues such as cerebral spinal fluid (CSF), the skull and fat tissue. Images are then affine registered using a 12-parameter transformation, where the optimisation is extended to include translations, rotations, uniaxial strains, and shears.

#### Nonlinear registration

Nonlinear registration requires the calculation of a three-dimensional displacement field at each voxel that maps the moving image to the target. The affine registered images are first smoothed by convolving with an isotropic Gaussian kernel with standard deviation σ=32mm, corresponding to full width half maximum (FWHM) of 75.3542. The cost function to be minimised expresses the intensity difference between the moving and target image and includes regularisation parameters which ensure a smooth displacement field and penalise large deformations. By design, the minimum of the cost function is found when the displacement field results in a good match between the moving and target images. Solving for the global minimum is made efficient by using the biconjugate gradients stabilised method. The displacement field, initially set as zero, is then updated. This is repeated three times, decreasing the smoothing of the images each time, σ is reduced to 16mm (FWHM=37.6771) then to 6mm (FWHM=14.1289), increasing the precision of the registration. The last iteration outputs a net displacement field. Finally, the rigid body, affine and nonlinear transformations are combined in a single transformation to be applied in a single interpolation step.

### Registration target

An iterative procedure was used to generate an atlas target consistent with the registration procedure. The initial registration target was chosen the T1w atlas image used by Hagler et al. (2009) for the AtlasTrack method. Participants’ T1w scans were registered to the T1w atlas image and the warps were applied to the eleven channels needed for the multimodal registration. A group average was computed for each channel across the 1000 scans with the best alignment with the registration target, based on the mean Pearson correlation across all eleven input channels. The registration target was updated with the new multimodal average and the procedure was repeated three times, producing an iteratively refined multimodal atlas. The group average was restricted to the 1000 participants most correlated to the registration as it has been shown that not all participants should be weighted equally (Wu et al., 2011) when constructing a group average. Up-weighting participants close to the population mean, while down-weighting others, will produce a sharper registration target and improve registration.

### Registration evaluation

#### Comparison with other registration algorithms

Our registration method was evaluated against two widely used registration algorithms. FSL’s FLIRT and FNIRT (from herein referred to only as “FSL”) and ANTs were applied to a random subset of 200 participants. The demographics of the evaluation subset are summarised in Table 2. A subset of only 200 was chosen because of the long computation time needed for ANTs. As ANTs is a multimodal registration algorithm and it too requires modalities to be mutually aligned prior to multimodal registration, the exact same input images and registration target used for MINT were used for ANTs. ANTs allows the inputs to each registration step (rigid body, affine, nonlinear) to vary, therefore the inputs for each step were matched to those of MINT. The registration target for ANTs was set to be the same as the final registration target in MINT to remove discrepancy in performance between the algorithms due to differences in registration targets. Given that FSL is a unimodal registration algorithm, the input images were each participant’s T1w scans and the registration target was the T1 channel from the final registration target in MINT.

**Table 2.** Summary demographics of the sample of 200 random observations used in registration evaluation

|  | Baseline | Two-year FU |
| --- | --- | --- |
| n | 134 | 66 |
| Age, median years (range) | 9.9 (8.9-10.9) | 12 (10.9-13) |
| Male sex, n (%) | 72 (53.7) | 36 (54.5) |
| *Parental education* | n (%) | n (%) |
| < HS diploma | 6 (4.5) | 2 (3.0) |
| HS diploma/GED | 6 (4.5) | 3 (4.5) |
| Some college | 17 (12.7) | 3 (4.5) |
| Bachelor degree | 33 (24.6) | 15 (22.7) |
| Postgraduate degree | 72 (53.7) | 43 (65.2) |
| NA | 0 (0) | 0 (0) |
| *Race (race.6level)* | n (%) | n (%) |
| AIAN/NHPI | 1 (0.75) | 1 (1.5) |
| Asian | 29 (21.6) | 17 (25.6) |
| Black | 2 (1.5) | 1 (1.5) |
| Mixed | 25 (18.6) | 11 (16.7) |
| Other | 12 (9.0) | 3 (4.5) |
| White | 59 (44.0) | 33 (50.0) |
| NA | 6 (4.5) | 0 (0) |
| *Hispanic ethnicity* | n (%) | n (%) |
| Yes | 32 (23.9) | 11 (16.7) |
| No | 100 (74.6) | 54 (81.8) |
| NA | 2 (1.5) | 1 (1.5) |
| *Household income* | n (%) | n (%) |
| <50k | 23 (17.2) | 3 (4.6) |
| 50-100k | 12 (9.0) | 10 (15.2) |
| >100k | 86 (64.2) | 48 (72.7) |
| NA | 13 (9.7) | 5 (7.6) |

#### Similarity measure

Numerous similarity measures have been employed in previous registration studies, primarily Dice or Jaccard overlap measures or measures based on mutual information calculations (Bhatia et al., 2007). However, these are pairwise assessments and do not describe the overall alignment across all participants. To determine the overall agreement in anatomical structures across participants, we can calculate the probability that a voxel is assigned to a given ROI. When there is good alignment between participants, voxel probabilities will be high within the ROI and drop off sharply; poor alignment will result in a blurry ROI with a gradual decrease in voxel probabilities. A blurrier ROI reflects greater uncertainty in the location of the structure’s boundary. Shannon entropy is a natural measure of uncertainty (Namdari and Li, 2019), based on the probability of an event occurring, and as such is our chosen similarity measure.

Following registration with each of the three algorithms tested, each participant’s subcortical grey matter ROIs and white matter ROIs from the Freesurfer parcellation and AtlasTrack, respectively, were transformed to the ABCD atlas space. ROIs were binarised and summed across participants and divided by the number of participants to produce a probabilistic ROI for each structure, according to each registration algorithm. Each voxel, *i* = 1, … *n*, then contains a probability that it belongs to a given anatomical structure. The Shannon entropy for each ROI was calculated as

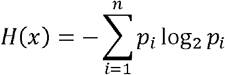

### Computation time

Given the importance of reasonable computation time for the spatial normalisation, we report the average time taken to register a single participant to the atlas.

### Incorporating multiple brain parcellations into the MINT ABCD atlas space

We incorporated two independent brain parcellations into the MINT ABCD space to provide parcellations of subcortical structures not available from Freesurfer or AtlasTrack. Different parcellation schemes offer varying degrees of detail with respect to different anatomical structures. This allows us to make inferences at different granularity levels and improve reproducibility across studies using different parcellation schemes. Therefore we included a subcortical atlas that was generated using T1 and T2 scans from 168 typical adults from the Human Connectome Project (HCP) (Pauli et al., 2018), and a thalamic nuclei atlas that was generated using a k-means algorithm taking as inputs the mean FOD SH coefficients from within a Freesurfer parcellation of the thalamus, using adult HCP data from 70 participants (Najdenovska et al., 2018).

### Application of the MINT ABCD atlas to statistical analyses

Fan et al. (2022) recently published the fast and efficient mixed-effects algorithm (FEMA) which implements linear mixed effects models for use in large-scale neuroimaging data. FEMA can perform whole-brain, mass univariate analyses on complex large-scale imaging data in a computationally efficient manner. This requires voxelwise correspondence across participants prior to statistical analyses; therefore, we demonstrate the use of the MINT atlas in conjunction with FEMA.

Associations between body mass index (BMI) and dMRI measures in adolescents from the ABCD study have previously been reported (Rapuano et al., 2020). Rapuano et al. demonstrated that greater BMI was associated with greater restricted normalised isotropic (RNI) diffusion in the intracellular tissue compartment, as measured by RSI within the nucleus accumbens, putamen, caudate, pallidum, ventral diencephalon, thalamus, amygdala and hippocampus. The study presented results from a region-of-interest (ROI) analysis, with a post hoc voxelwise analysis restricted only to voxels within the ROIs studied. Here we replicate and expand upon these results to demonstrate the validity of voxelwise analysis following registration using MINT.

After spatial normalisation using MINT, we measured voxelwise associations between BMI and RNI across the whole brain. RNI is equivalent to the restricted isotropic component in the study by Rapuano et al (2020), also referred to as cellular density. Here we report our findings using the updated nomenclature from ABCD Study Data Release 4.0 (release notes with further details are available from https://nda.nih.gov/abcd). We used FEMA to estimate univariate general linear mixed effects models at each voxel to test the associations between BMI and RNI. In our model we included household income, parental education, Hispanic ethnicity, self-declared race, parental marital status, intracranial volume, scanner ID and MRI software version as fixed effects and participant ID and family relatedness as random effects. Covariates were chosen to match those included by Rapuano et al., (2020). The demographics of the sample used in this analysis are summarised in Table 3.

**Table 3.** Summary demographics of participants analysed in the statistical analysis application of the MINT atlas.

|  | Baseline |
| --- | --- |
| n | 8247 |
| Age, median years (range) | 9.9 (8.9-11) |
| Male sex, n (%) | 4224 (51.2) |
| BMI, median kg/m<sup>2</sup> (range) | 17.5 (11.1-35.7) |
| *Parental education* | n (%) |
| < HS diploma | 262 (3.2) |
| HS diploma/GED | 606 (7.3) |
| Some college | 2066 (25.1) |
| Bachelor degree | 2267 (27.5) |
| Postgraduate degree | 3046 (36.9) |
| NA | 0 (0) |
| *Race (race.6level)* | n (%) |
| AIAN/NHPI | 49 (0.6) |
| Asian | 170 (2.1) |
| Black | 1034 (12.5) |
| Mixed | 1016 (12.3) |
| Other | 328 (4.0) |
| White | 5650 (68.5) |
| NA | 0 (0) |
| Hispanic ethnicity, n (%) | 1593 (19.3) |
| *Household income* | n (%) |
| <50k | 2195 (26.6) |
| 50-100k | 2365 (28.7) |
| >100k | 3687 (44.7) |
| NA | 0 (0) |

In addition to studying scalar indices such as RNI, nonlinear registration allows us to examine voxelwise local tissue contraction or expansion relative to the atlas and its association with our variable of interest, which is not possible with ROI analyses. From the nonlinear spatial transformations that warp each participant’s affine registered images to the atlas, we calculated the determinant of the spatial derivative matrix, the Jacobian, which describes the local tissue expansion or contraction. We measured voxelwise associations between BMI and the Jacobian (JA), including the same covariates in the model as described above.

## Results

### ABCD MINT atlas

Figure 1 shows the eleven channels of the multimodal ABCD MINT atlas. The T1w, white and grey matter segmentation channels (Figure 1a, b and c, respectively) show the preservation of anatomical detail in the cortical and subcortical structures. The diffusion channels of the zeroth order coefficients for the restricted, hindered, and free water compartments (Figure 1d, j, k) show distinct tissue properties. The diffusion channels of the second order SH coefficients for the restricted FOD (Figure 1e-j) show the orientational information contained in dMRI data, particularly in the white matter, which help drive alignment across participants in oriented structures. Figure 2 shows the average Freesurfer parcellation, the color-coded FA map, and the restricted and hindered fourth order FODs from the ABCD atlas. Table 4 describes the demographics of the 1000 subjects used to create the final ABCD atlas.

**Table 4.** Summary demographics of the 1000 participants included in the final MINT atlas.

|  | Baseline | Two-year FU |
| --- | --- | --- |
| n | 608 | 391 |
| Age, median years (range) | 9.9 (8.9-11) | 11.8 (10.8-13.5) |
| Male sex, n (%) | 306 (50.3) | 200 (51.2) |
| *Parental education* | n (%) | n (%) |
| < HS diploma | 64 (10.5) | 40 (10.2) |
| HS diploma/GED | 57 (9.4) | 38 (9.7) |
| Some college | 133 (21.9) | 88 (22.5) |
| Bachelor degree | 138 (22.7) | 83 (21.2) |
| Postgraduate degree | 215 (35.4) | 142 (36.3) |
| NA | 1 (0.2) | 0 (0) |
| *Race (race.6level)* | n (%) | n (%) |
| AIAN/NHPI | 7 (1.2) | 6 (1.5) |
| Asian | 72 (11.8) | 41 (10.5) |
| Black | 19 (3.1) | 11 (2.8) |
| Mixed | 100 (16.4) | 52 (13.3) |
| Other | 87 (14.3) | 57 (14.6) |
| White | 287 (47.2) | 207 (52.9) |
| NA | 36 (5.9) | 17 (4.3) |
| *Hispanic ethnicity* | n (%) | n (%) |
| Yes | 265 (43.6) | 175 (44.8) |
| No | 329 (54.1) | 211 (54.0) |
| NA | 14 (2.3) | 5 (1.2) |
| *Household income* | n (%) | n (%) |
| <50k | 176 (28.9) | 105 (26.9) |
| 50-100k | 103 (16.9) | 81 (20.7) |
| >100k | 259 (42.6) | 171 (43.7) |
| NA | 70 (11.5) | 34 (8.7) |

**Figure 1.**
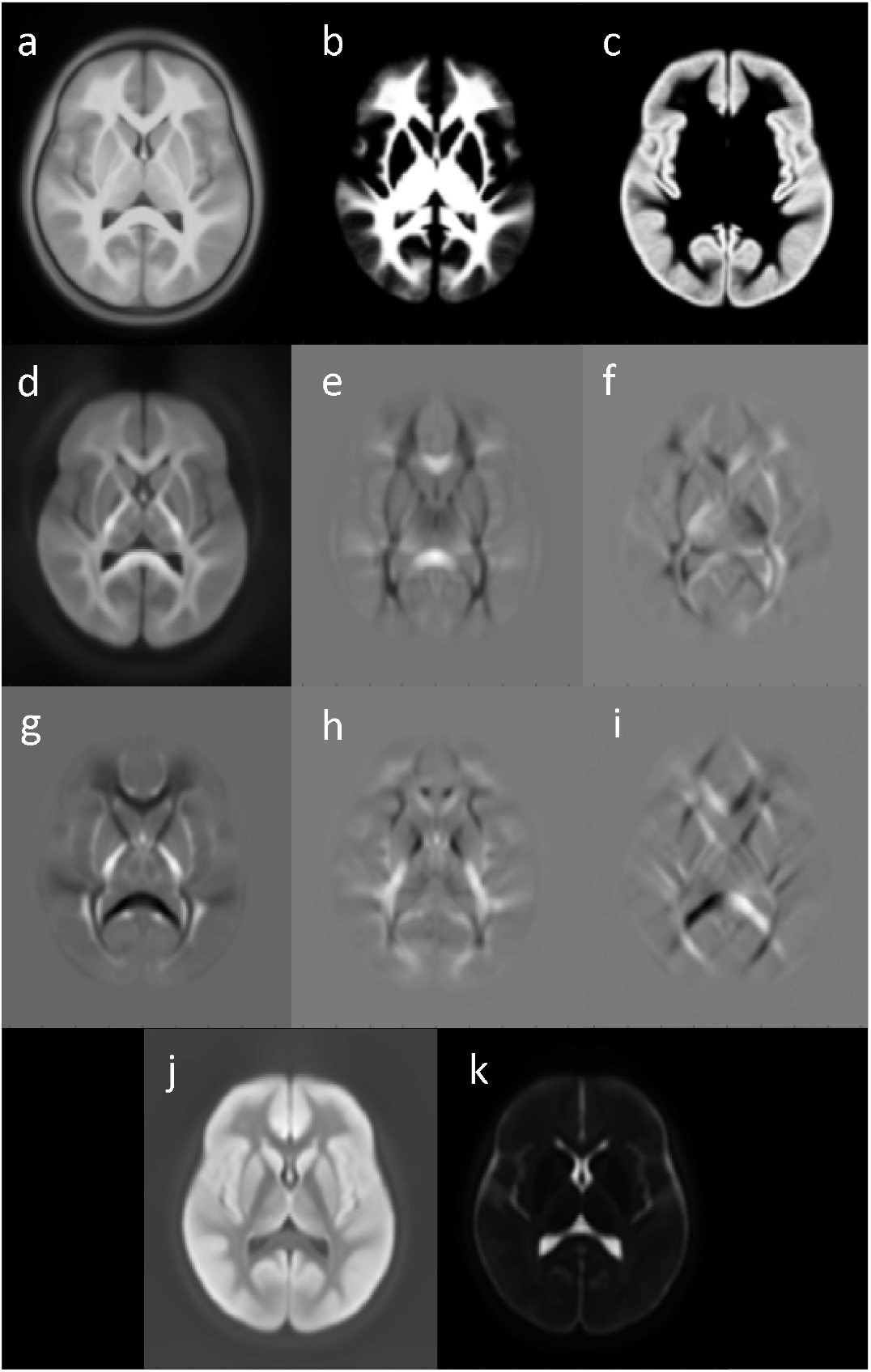
Eleven channels of the MINT ABCDatlas: (a) T1w; (b) white matter segmentation; (c) grey matter segmentation; (d) zeroth order spherical harmonics coefficient of the restricted FOD; (e-i) second order, -2 to 2 degree spherical harmonics coefficients of the restricted FOD; (j) zeroth order spherical harmonics coefficient of the hindered FOD; (k) zeroth order spherical harmonics coefficient of the free water FOD.

**Figure 2.**
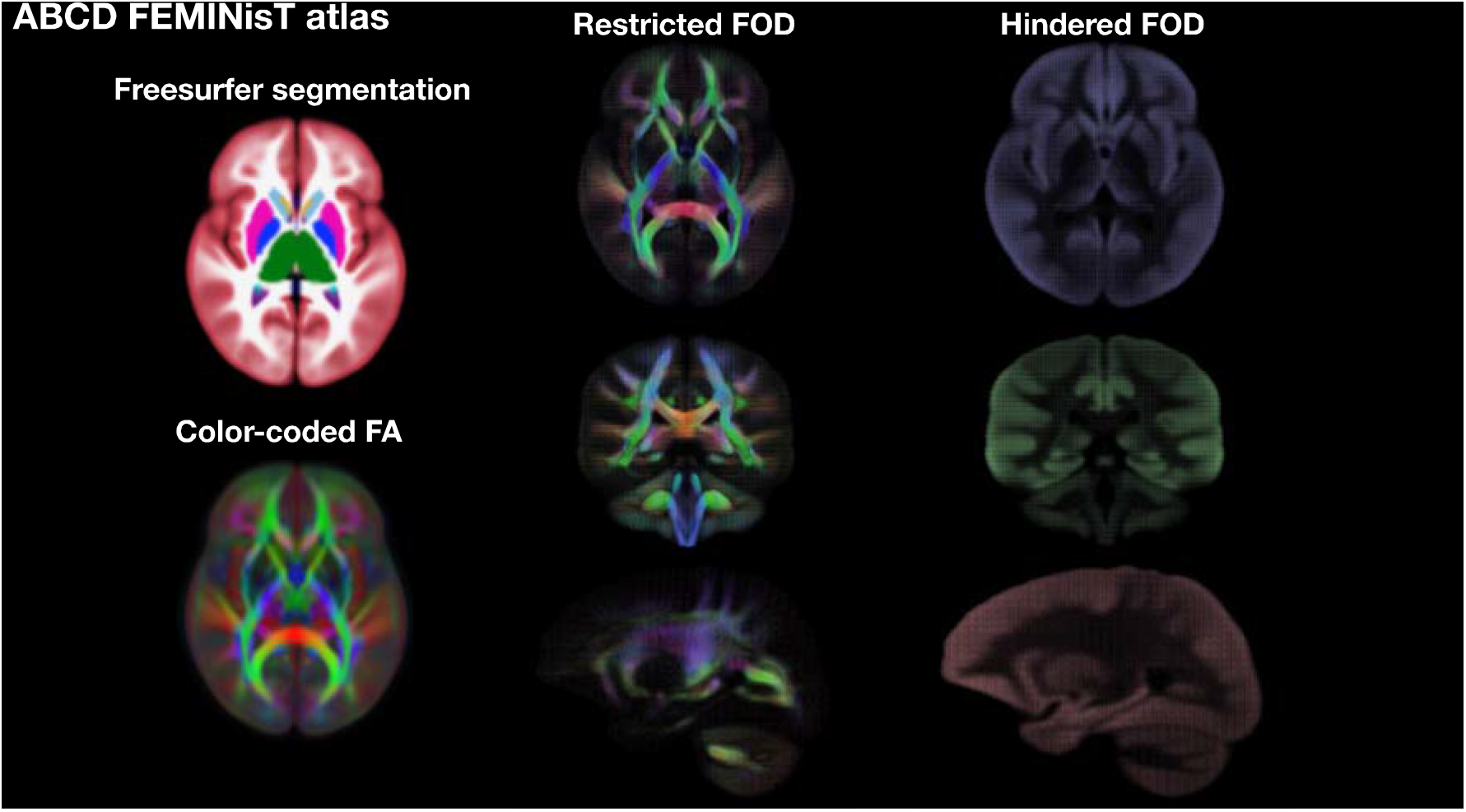
The MINT ABCDatlas maps for the Freesurfer tissue segmentation, color-coded FA, restricted and hindered FODs.

### Registration evaluation

We evaluated the performance of MINT against FSL and ANTs. Figure 3 shows the entropy calculations for white matter and subcortical grey matter ROIs for MINT, ANTs and FSL. Since entropy is a measure of uncertainty, lower entropy values represent greater certainty, therefore a sharper ROI owing to better alignment across participants.

**Figure 3.**
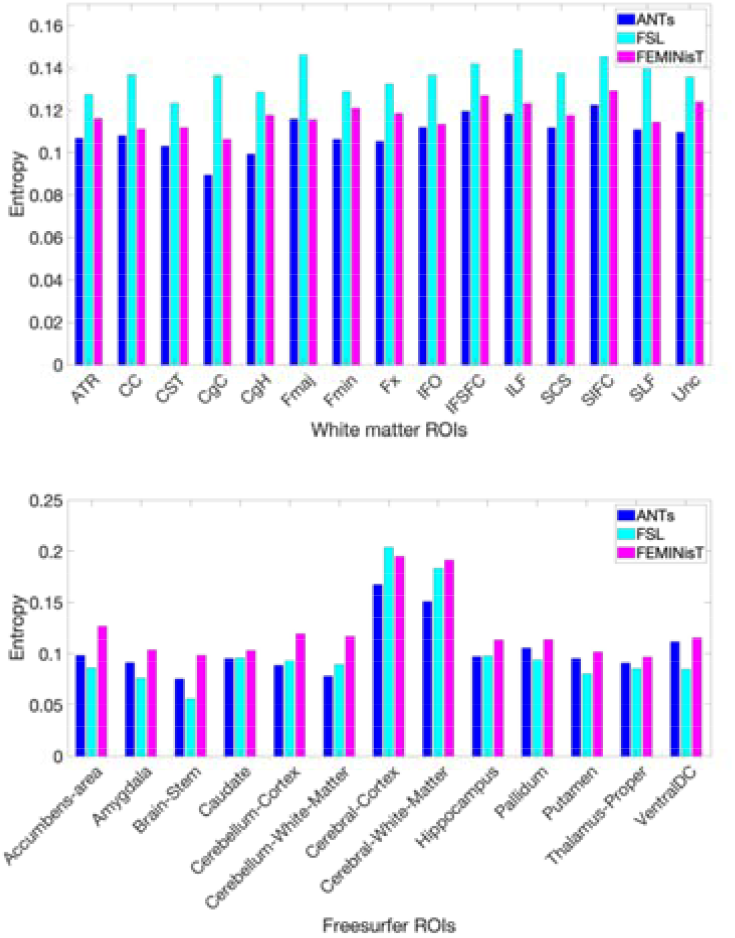
ROI entropy measures for each registration algorithm for white matter ROIs from AtlasTrack and tissue segmentation ROIs from Freesurfer.

MINT and ANTs show comparable performances in alignment of white matter tracts, whereas FSL-aligned tracts have markedly higher entropy values. Figure 4 shows the voxelwise entropy measure for the corpus callosum produced by each registration algorithm. In the alignment of subcortical grey matter, overall FSL performs better than both ANTs and MINT (Figure 3), as illustrated by slightly sharper voxelwise entropy maps of the putamen (Figure 4).

**Figure 4.**
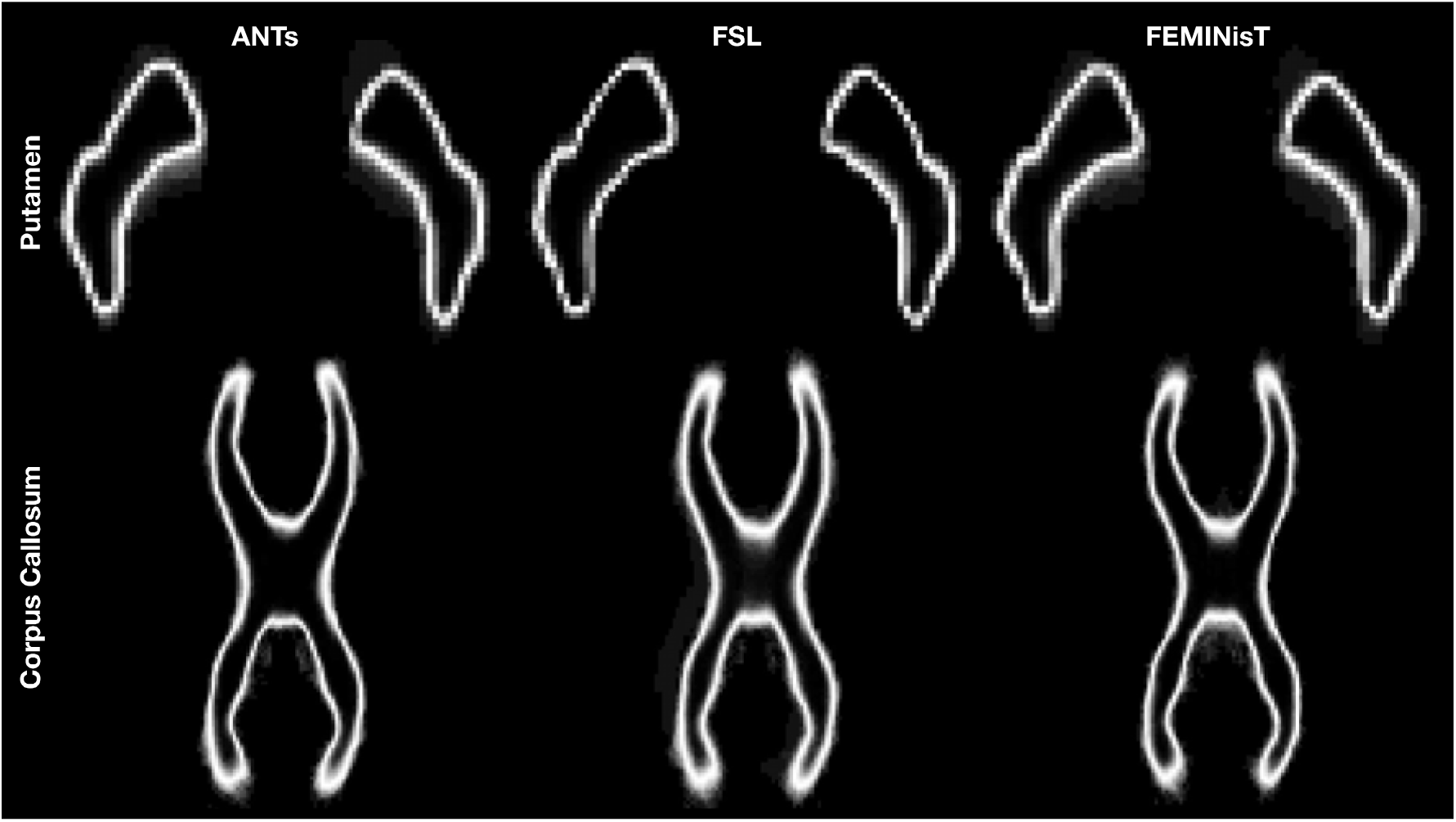
Voxelwise entropy measure for each registration algorithm for the putamen and corpus callosum ROIs.

### Computation time

The average computation time for registering a single participant to the atlas was 29 hrs 1 min 41sec for ANTs, 30 min 24 sec for FSL and 24 min 35 sec for MINT.

### Application of the MINT ABCD atlas to statistical analyses

Using the MINT ABCD atlas we analysed whole brain, voxelwise associations between BMI and RNI and JA. Figure 5 shows greater BMI was associated with greater RNI across the brain, most notably in the subcortical grey matter, including the nucleus accumbens, caudate, globus pallidus, putamen and thalamus, in agreement with the findings of Rapuano et al. (Rapuano et al., 2020). Form our voxelwise analysis we see that effects are distributed nonuniformly across and within ROIs, highlighting the advantages of voxelwise analyses over ROI-based approaches. Furthermore, using a whole-brain approach instead of ROI-based, we can see that significant associations are not limited to these subcortical structures. There are significant, positive associations between BMI and RNI in white matter surrounding the subcortical grey matter, the forceps minor, frontal white matter, and the cerebellum.

**Figure 5.**
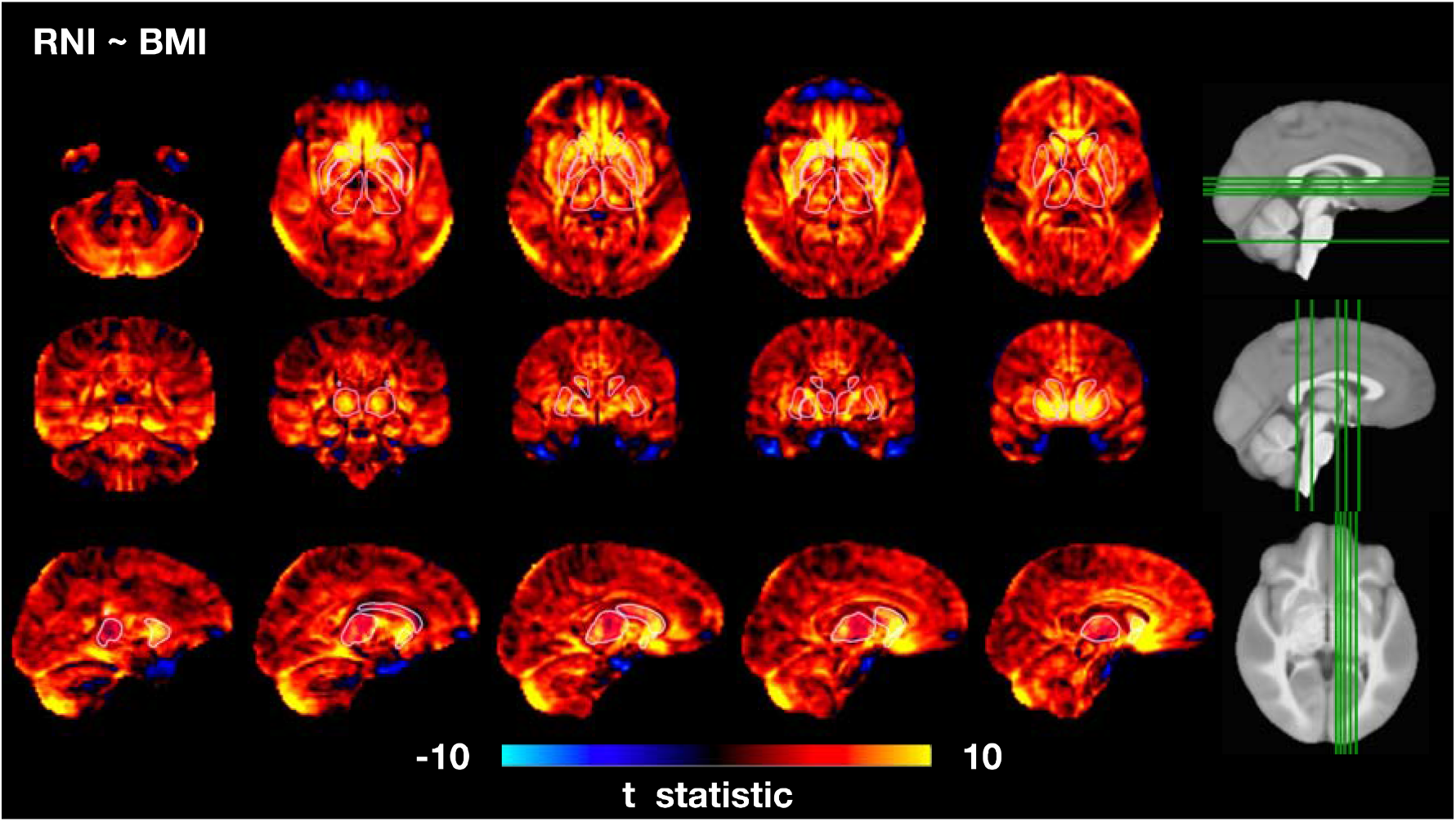
Voxelwise associations between RNI and BMI. ROI outlines are shown for the putamen, caudate, nucleus accumbens, pallidum and the thalamus.

Using the JA output by MINT, we investigated the associations between BMI and localised brain tissue volume differences. Figure 6 shows that greater BMI is associated with larger volume in the pallidum, centrum semiovale, periventricular white matter, the white matter surrounding the thalamus and putamen, and cerebellar white matter. Greater BMI was associated with smaller volume in the head of the caudate, the putamen, nucleus accumbens, and cerebellar cortex. Associations within the thalamus varied, with negative associations in the anterior dorsal nuclei and positive associations in the posterior ventral nuclei. Similarly, associations with the corpus callosum were spatially variable. The splenium of the corpus callosum showed positive associations between BMI and JA, while the rest of the corpus callosum showed negative associations.

**Figure 6.**
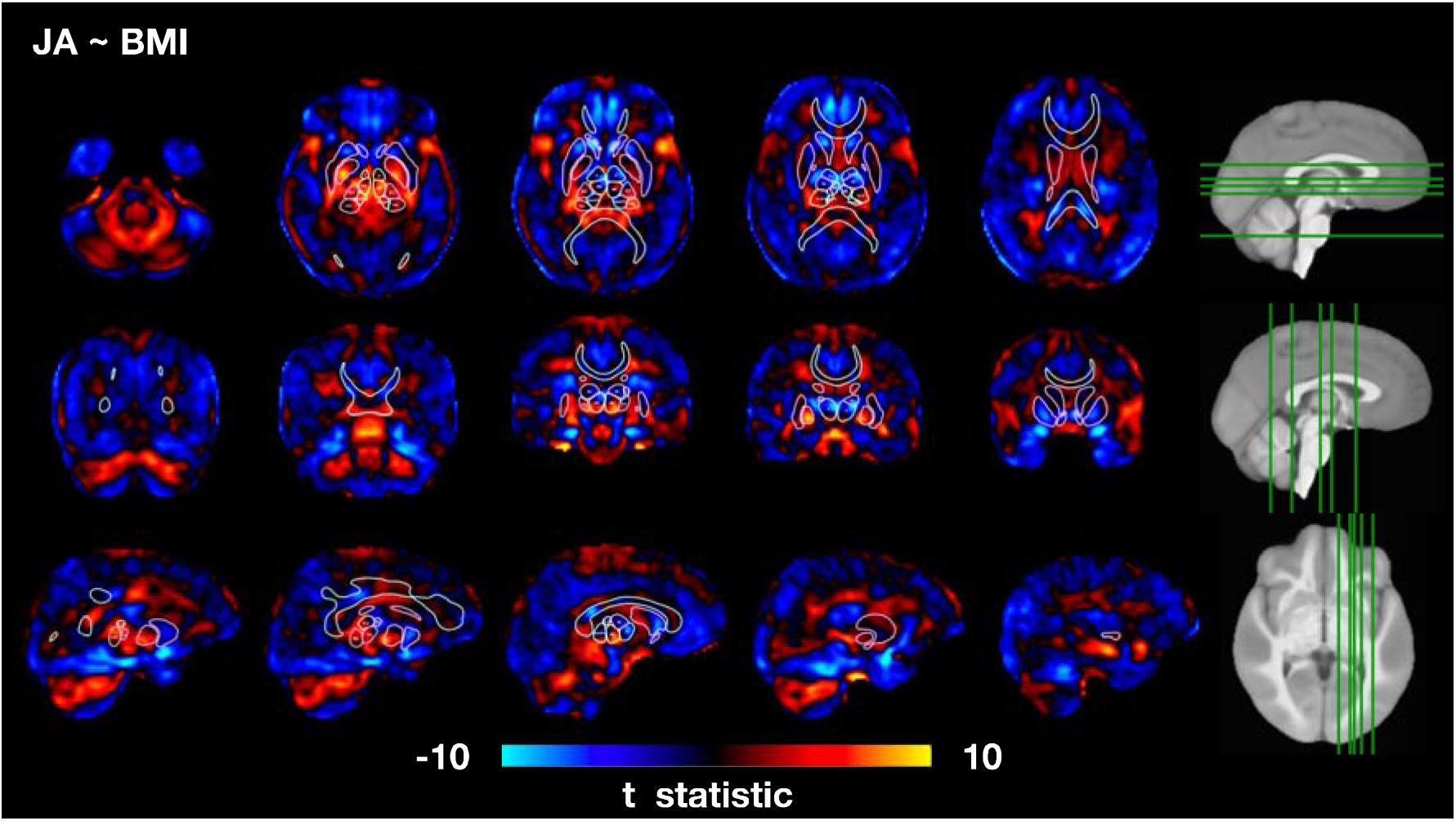
Voxelwise associations between the Jacobians (JA) from the MINT registration and BMI. ROI outlines are shown for the corpus callosum, putamen, caudate, nucleus accumbens, pallidum and the thalamic nuclei. The thalamic nuclei are: VA=ventral anterior, A=anterior, VA=ventral, MD=mediodorsal, VLV=ventral-latero-ventral, VLD=ventral-latero-dorsal, clpmP=central-latero-lateral posterior-medial-pulvinar, PUL=pulvinar.

## Discussion

We have presented a publicly available multimodal atlas constructed from ABCD data. Using the multimodal registration algorithm from MINT we can align scans from thousands of participants in eleven modalities simultaneously, achieving comparable performance to current state-of-the-art multimodal registration, at a fraction of the computation time.

There were some differences in performance for the three registration algorithms across the different tissue types. Overall, the registration accuracy of MINT was comparable to that of ANTs in white and subcortical grey matter. Both ANTs and MINT were outperformed by FSL in the alignment of certain subcortical ROIs. Both ANTs and MINT require within-participant alignment of scans across modalities prior to alignment of scans across participants. It may be that this extra step introduces small misalignments which could be propagated to the between-participant registration step. Furthermore, the boundaries of subcortical structures are best observed on T1w scans, and it is possible that this is sufficient for good alignment of these structures. Conversely, the two multimodal produced substantially better alignment across white matter tracts than FSL. The orientational information in white matter is captured by the eight dMRI channels improving alignment in these structures, while the T1w channel is not sensitive to this information, and therefore performs worse.

Using the ABCD MINT atlas and FEMA we have demonstrated that greater BMI was associated with greater RNI across the brain, reproducing the findings from Rapuano et al. (2020). The agreement between our results and those previously published validates the use of MINT-registered data for whole-brain voxelwise analysis. We also presented findings of associations between BMI and JA that would not have been possible to uncover using a ROI approach. We demonstrated spatially varying associations between BMI and both RNI and JA. An in-depth study of these results is beyond the scope of this paper; however, they demonstrate the utility of MINT and FEMA.

The inclusion of the atlas of thalamic nuclei (Najdenovska et al., 2018) allowed us to identify the distinct thalamic nuclei in the varying associations between JA and BMI. Incorporating external, publicly available parcellations into the MINT ABCD space provided greater detail to our analysis than was possible using solely the Freesurfer segmentation. Multiple segmentation schemes within the same space should lead to improved reproducibility across studies by reducing the sources of method variation between studies. This is particularly important in large, densely phenotyped studies such as ABCD where many investigators will want to query the same data but using different approaches.

Moreover, brain morphology has been shown to exhibit variation across ancestral groups (Bakken et al., 2011; Fan et al., 2015) and registration accuracy is approved when the atlas is representative of the study cohort (Xie et al., 2015; Zhao et al., 2019). The size and demographic diversity of the ABCD sample used to generate the atlas make it applicable to other paediatric studies, enabling more reliable comparisons across studies.

In conclusion, we have presented the ABCD MINT atlas as a publicly available resource to facilitate whole brain voxelwise analyses for the ABCD study. Our multimodal registration algorithm allows us to incorporate multiple imaging modalities, with good spatial alignment across participants, while keeping computation time at a minimum, ensuring that this approach is scalable to include the future timepoints from this longitudinal study.

## Acknowledgements

The authors wish to thank the youth and families participating in the Adolescent Brain Cognitive Development (ABCD) Study and all ABCD staff involved in data collection and curation. Dr. Dale reports that he was a Founder of and holds equity in CorTechs Labs, Inc., and serves on its Scientific Advisory Board. He is a member of the Scientific Advisory Board of Human Longevity, Inc. He receives funding through research grants from GE Healthcare to UCSD. The terms of these arrangements have been reviewed by and approved by UCSD in accordance with its conflict of interest policies.

## References

Andersson, J.L., Skare, S., Ashburner, J., 2003. How to correct susceptibility distortions in spin-echo echo-planar images: application to diffusion tensor imaging. Neuroimage 20, 870–888.

Avants, B.B., Duda, J.T., Kilroy, E., Krasileva, K., Jann, K., Kandel, B.T., Tustison, N.J., Yan, L., Jog, M., Smith, R., Wang, Y., Dapretto, M., Wang, D.J., 2015. The pediatric template of brain perfusion. Sci Data 2, 150003.

Avants, B.B., Epstein, C.L., Grossman, M., Gee, J.C., 2008. Symmetric diffeomorphic image registration with cross-correlation: evaluating automated labeling of elderly and neurodegenerative brain. Med Image Anal 12, 26–41.

Avants, B.B., Tustison, N.J., Stauffer, M., Song, G., Wu, B., Gee, J.C., 2014. The Insight ToolKit image registration framework. Front Neuroinform 8, 44.

Bakken, T.E., Dale, A.M., Schork, N.J., 2011. A geographic cline of skull and brain morphology among individuals of European Ancestry. Hum Hered 72, 35–44.

Ball, G., Malpas, C.B., Genc, S., Efron, D., Sciberras, E., Anderson, V., Nicholson, J.M., Silk, T.J., 2018. Multimodal Structural Neuroimaging Markers of Brain Development and ADHD Symptoms. Am J Psychiatry, appiajp201818010034.

Bethlehem, R.A.I., Seidlitz, J., White, S.R., Vogel, J.W., Anderson, K.M., Adamson, C., Adler, S., Alexopoulos, G.S., Anagnostou, E., Areces-Gonzalez, A., Astle, D.E., Auyeung, B., Ayub, M., Bae, J., Ball, G., Baron-Cohen, S., Beare, R., Bedford, S.A., Benegal, V., Beyer, F., Blangero, J., Blesa Cabez, M., Boardman, J.P., Borzage, M., Bosch-Bayard, J.F., Bourke, N., Calhoun, V.D., Chakravarty, M.M., Chen, C., Chertavian, C., Chetelat, G., Chong, Y.S., Cole, J.H., Corvin, A., Costantino, M., Courchesne, E., Crivello, F., Cropley, V.L., Crosbie, J., Crossley, N., Delarue, M., Delorme, R., Desrivieres, S., Devenyi, G.A., Di Biase, M.A., Dolan, R., Donald, K.A., Donohoe, G., Dunlop, K., Edwards, A.D., Elison, J.T., Ellis, C.T., Elman, J.A., Eyler, L., Fair, D.A., Feczko, E., Fletcher, P.C., Fonagy, P., Franz, C.E., Galan-Garcia, L., Gholipour, A., Giedd, J., Gilmore, J.H., Glahn, D.C., Goodyer, I.M., Grant, P.E., Groenewold, N.A., Gunning, F.M., Gur, R.E., Gur, R.C., Hammill, C.F., Hansson, O., Hedden, T., Heinz, A., Henson, R.N., Heuer, K., Hoare, J., Holla, B., Holmes, A.J., Holt, R., Huang, H., Im, K., Ipser, J., Jack, C.R., Jr., Jackowski, A.P., Jia, T., Johnson, K.A., Jones, P.B., Jones, D.T., Kahn, R.S., Karlsson, H., Karlsson, L., Kawashima, R., Kelley, E.A., Kern, S., Kim, K.W., Kitzbichler, M.G., Kremen, W.S., Lalonde, F., Landeau, B., Lee, S., Lerch, J., Lewis, J.D., Li, J., Liao, W., Liston, C., Lombardo, M.V., Lv, J., Lynch, C., Mallard, T.T., Marcelis, M., Markello, R.D., Mathias, S.R., Mazoyer, B., McGuire, P., Meaney, M.J., Mechelli, A., Medic, N., Misic, B., Morgan, S.E., Mothersill, D., Nigg, J., Ong, M.Q.W., Ortinau, C., Ossenkoppele, R., Ouyang, M., Palaniyappan, L., Paly, L., Pan, P.M., Pantelis, C., Park, M.M., Paus, T., Pausova, Z., Paz-Linares, D., Pichet Binette, A., Pierce, K., Qian, X., Qiu, J., Qiu, A., Raznahan, A., Rittman, T., Rodrigue, A., Rollins, C.K., Romero-Garcia, R., Ronan, L., Rosenberg, M.D., Rowitch, D.H., Salum, G.A., Satterthwaite, T.D., Schaare, H.L., Schachar, R.J., Schultz, A.P., Schumann, G., Scholl, M., Sharp, D., Shinohara, R.T., Skoog, I., Smyser, C.D., Sperling, R.A., Stein, D.J., Stolicyn, A., Suckling, J., Sullivan, G., Taki, Y., Thyreau, B., Toro, R., Traut, N., Tsvetanov, K.A., Turk-Browne, N.B., Tuulari, J.J., Tzourio, C., Vachon-Presseau, E., Valdes-Sosa, M.J., Valdes-Sosa, P.A., Valk, S.L., van Amelsvoort, T., Vandekar, S.N., Vasung, L., Victoria, L.W., Villeneuve, S., Villringer, A., Vertes, P.E., Wagstyl, K., Wang, Y.S., Warfield, S.K., Warrier, V., Westman, E., Westwater, M.L., Whalley, H.C., Witte, A.V., Yang, N., Yeo, B., Yun, H., Zalesky, A., Zar, H.J., Zettergren, A., Zhou, J.H., Ziauddeen, H., Zugman, A., Zuo, X.N., R, B., Aibl, Alzheimer’s Disease Neuroimaging, I., Alzheimer’s Disease Repository Without Borders, I., Team, C., Cam, C.A.N., Ccnp, Cobre, cVeda, Group, E.D.B.A.W., Developing Human Connectome, P., FinnBrain, Harvard Aging Brain, S., Imagen, Kne, Mayo Clinic Study of, A., Nspn, Pond, Group, P.-A.R., Vetsa, Bullmore, E.T., Alexander-Bloch, A.F., 2022. Brain charts for the human lifespan. Nature 604, 525–533.

Bhatia, K.K., Hajnal, J., Hammers, A., Rueckert, D., 2007. Similarity metrics for groupwise non-rigid registration. Med Image Comput Comput Assist Interv 10, 544–552.

Bos, M.G.N., Wierenga, L.M., Blankenstein, N.E., Schreuders, E., Tamnes, C.K., Crone, E.A., 2018. Longitudinal structural brain development and externalizing behavior in adolescence. J Child Psychol Psychiatry 59, 1061–1072.

Brown, T.T., Kuperman, J.M., Chung, Y., Erhart, M., McCabe, C., Hagler, D.J., Jr., Venkatraman, V.K., Akshoomoff, N., Amaral, D.G., Bloss, C.S., Casey, B.J., Chang, L., Ernst, T.M., Frazier, J.A., Gruen, J.R., Kaufmann, W.E., Kenet, T., Kennedy, D.N., Murray, S.S., Sowell, E.R., Jernigan, T.L., Dale, A.M., 2012. Neuroanatomical assessment of biological maturity. Curr Biol 22, 1693–1698.

Brunsing, R.L., Schenker-Ahmed, N.M., White, N.S., Parsons, J.K., Kane, C., Kuperman, J., Bartsch, H., Kader, A.K., Rakow-Penner, R., Seibert, T.M., Margolis, D., Raman, S.S., McDonald, C.R., Farid, N., Kesari, S., Hansel, D., Shabaik, A., Dale, A.M., Karow, D.S., 2017. Restriction spectrum imaging: An evolving imaging biomarker in prostate MRI. J Magn Reson Imaging 45, 323–336.

Casey, B.J., Cannonier, T., Conley, M.I., Cohen, A.O., Barch, D.M., Heitzeg, M.M., Soules, M.E., Teslovich, T., Dellarco, D.V., Garavan, H., Orr, C.A., Wager, T.D., Banich, M.T., Speer, N.K., Sutherland, M.T., Riedel, M.C., Dick, A.S., Bjork, J.M., Thomas, K.M., Chaarani, B., Mejia, M.H., Hagler, D.J., Jr., Daniela Cornejo, M., Sicat, C.S., Harms, M.P., Dosenbach, N.U.F., Rosenberg, M., Earl, E., Bartsch, H., Watts, R., Polimeni, J.R., Kuperman, J.M., Fair, D.A., Dale, A.M., Workgroup, A.I.A., 2018. The Adolescent Brain Cognitive Development (ABCD) study: Imaging acquisition across 21 sites. Dev Cogn Neurosci 32, 43–54.

Casey, B.J., Heller, A.S., Gee, D.G., Cohen, A.O., 2019. Development of the emotional brain. Neurosci Lett 693, 29–34.

Casey, B.J., Tottenham, N., Liston, C., Durston, S., 2005. Imaging the developing brain: what have we learned about cognitive development? Trends Cogn Sci 9, 104–110.

Dick, A.S., Lopez, D.A., Watts, A.L., Heeringa, S., Reuter, C., Bartsch, H., Fan, C.C., Kennedy, D.N., Palmer, C., Marshall, A., Haist, F., Hawes, S., Nichols, T.E., Barch, D.M., Jernigan, T.L., Garavan, H., Grant, S., Pariyadath, V., Hoffman, E., Neale, M., Stuart, E.A., Paulus, M.P., Sher, K.J., Thompson, W.K., 2021. Meaningful associations in the adolescent brain cognitive development study. Neuroimage 239, 118262.

Fan, C.C., Bartsch, H., Schork, A.J., Chen, C.H., Wang, Y., Lo, M.T., Brown, T.T., Kuperman, J.M., Hagler, D.J., Jr., Schork, N.J., Jernigan, T.L., Dale, A.M., Pediatric Imaging, N., Genetics, S., 2015. Modeling the 3D geometry of the cortical surface with genetic ancestry. Curr Biol 25, 1988–1992.

Fischl, B., Salat, D.H., Busa, E., Albert, M., Dieterich, M., Haselgrove, C., van der Kouwe, A., Killiany, R., Kennedy, D., Klaveness, S., Montillo, A., Makris, N., Rosen, B., Dale, A.M., 2002. Whole brain segmentation: automated labeling of neuroanatomical structures in the human brain. Neuron 33, 341–355.

Fjell, A.M., Grydeland, H., Krogsrud, S.K., Amlien, I., Rohani, D.A., Ferschmann, L., Storsve, A.B., Tamnes, C.K., Sala-Llonch, R., Due-Tonnessen, P., Bjornerud, A., Solsnes, A.E., Haberg, A.K., Skranes, J., Bartsch, H., Chen, C.H., Thompson, W.K., Panizzon, M.S., Kremen, W.S., Dale, A.M., Walhovd, K.B., 2015. Development and aging of cortical thickness correspond to genetic organization patterns. Proc Natl Acad Sci U S A 112, 15462–15467.

Fonov, V., Evans, A.C., Botteron, K., Almli, C.R., McKinstry, R.C., Collins, D.L., Brain Development Cooperative, G., 2011. Unbiased average age-appropriate atlases for pediatric studies. Neuroimage 54, 313–327.

Garavan, H., Bartsch, H., Conway, K., Decastro, A., Goldstein, R.Z., Heeringa, S., Jernigan, T., Potter, A., Thompson, W., Zahs, D., 2018. Recruiting the ABCD sample: Design considerations and procedures. Dev Cogn Neurosci 32, 16–22.

Genc, S., Seal, M.L., Dhollander, T., Malpas, C.B., Hazell, P., Silk, T.J., 2017. White matter alterations at pubertal onset. Neuroimage 156, 286–292.

Hagler, D.J., Jr., Ahmadi, M.E., Kuperman, J., Holland, D., McDonald, C.R., Halgren, E., Dale, A.M., 2009. Automated white-matter tractography using a probabilistic diffusion tensor atlas: Application to temporal lobe epilepsy. Hum Brain Mapp 30, 1535–1547.

Hagler, D.J., Jr., Hatton, S., Cornejo, M.D., Makowski, C., Fair, D.A., Dick, A.S., Sutherland, M.T., Casey, B.J., Barch, D.M., Harms, M.P., Watts, R., Bjork, J.M., Garavan, H.P., Hilmer, L., Pung, C.J., Sicat, C.S., Kuperman, J., Bartsch, H., Xue, F., Heitzeg, M.M., Laird, A.R., Trinh, T.T., Gonzalez, R., Tapert, S.F., Riedel, M.C., Squeglia, L.M., Hyde, L.W., Rosenberg, M.D., Earl, E.A., Howlett, K.D., Baker, F.C., Soules, M., Diaz, J., de Leon, O.R., Thompson, W.K., Neale, M.C., Herting, M., Sowell, E.R., Alvarez, R.P., Hawes, S.W., Sanchez, M., Bodurka, J., Breslin, F.J., Morris, A.S., Paulus, M.P., Simmons, W.K., Polimeni, J.R., van der Kouwe, A., Nencka, A.S., Gray, K.M., Pierpaoli, C., Matochik, J.A., Noronha, A., Aklin, W.M., Conway, K., Glantz, M., Hoffman, E., Little, R., Lopez, M., Pariyadath, V., Weiss, S.R., Wolff-Hughes, D.L., DelCarmen-Wiggins, R., Feldstein Ewing, S.W., Miranda-Dominguez, O., Nagel, B.J., Perrone, A.J., Sturgeon, D.T., Goldstone, A., Pfefferbaum, A., Pohl, K.M., Prouty, D., Uban, K., Bookheimer, S.Y., Dapretto, M., Galvan, A., Bagot, K., Giedd, J., Infante, M.A., Jacobus, J., Patrick, K., Shilling, P.D., Desikan, R., Li, Y., Sugrue, L., Banich, M.T., Friedman, N., Hewitt, J.K., Hopfer, C., Sakai, J., Tanabe, J., Cottler, L.B., Nixon, S.J., Chang, L., Cloak, C., Ernst, T., Reeves, G., Kennedy, D.N., Heeringa, S., Peltier, S., Schulenberg, J., Sripada, C., Zucker, R.A., Iacono, W.G., Luciana, M., Calabro, F.J., Clark, D.B., Lewis, D.A., Luna, B., Schirda, C., Brima, T., Foxe, J.J., Freedman, E.G., Mruzek, D.W., Mason, M.J., Huber, R., McGlade, E., Prescot, A., Renshaw, P.F., Yurgelun-Todd, D.A., Allgaier, N.A., Dumas, J.A., Ivanova, M., Potter, A., Florsheim, P., Larson, C., Lisdahl, K., Charness, M.E., Fuemmeler, B., Hettema, J.M., Maes, H.H., Steinberg, J., Anokhin, A.P., Glaser, P., Heath, A.C., Madden, P.A., Baskin-Sommers, A., Constable, R.T., Grant, S.J., Dowling, G.J., Brown, S.A., Jernigan, T.L., Dale, A.M., 2019. Image processing and analysis methods for the Adolescent Brain Cognitive Development Study. Neuroimage 202, 116091.

Holland, D., Dale, A.M., Alzheimer’s Disease Neuroimaging, I., 2011. Nonlinear registration of longitudinal images and measurement of change in regions of interest. Med Image Anal 15, 489–497.

Joshi, S., Davis, B., Jomier, M., Gerig, G., 2004. Unbiased diffeomorphic atlas construction for computational anatomy. Neuroimage 23 Suppl 1, S151–160.

Jovicich, J., Czanner, S., Greve, D., Haley, E., van der Kouwe, A., Gollub, R., Kennedy, D., Schmitt, F., Brown, G., Macfall, J., Fischl, B., Dale, A., 2006. Reliability in multi-site structural MRI studies: effects of gradient non-linearity correction on phantom and human data. Neuroimage 30, 436–443.

Keihaninejad, S., Ryan, N.S., Malone, I.B., Modat, M., Cash, D., Ridgway, G.R., Zhang, H., Fox, N.C., Ourselin, S., 2012. The importance of group-wise registration in tract based spatial statistics study of neurodegeneration: a simulation study in Alzheimer’s disease. PLoS One 7, e45996.

Klein, A., Andersson, J., Ardekani, B.A., Ashburner, J., Avants, B., Chiang, M.C., Christensen, G.E., Collins, D.L., Gee, J., Hellier, P., Song, J.H., Jenkinson, M., Lepage, C., Rueckert, D., Thompson, P., Vercauteren, T., Woods, R.P., Mann, J.J., Parsey, R.V., 2009. Evaluation of 14 nonlinear deformation algorithms applied to human brain MRI registration. Neuroimage 46, 786–802.

Lebel, C., Beaulieu, C., 2011. Longitudinal development of human brain wiring continues from childhood into adulthood. J Neurosci 31, 10937–10947.

Lebel, C., Deoni, S., 2018. The development of brain white matter microstructure. Neuroimage.

Lenroot, R.K., Gogtay, N., Greenstein, D.K., Wells, E.M., Wallace, G.L., Clasen, L.S., Blumenthal, J.D., Lerch, J., Zijdenbos, A.P., Evans, A.C., Thompson, P.M., Giedd, J.N., 2007. Sexual dimorphism of brain developmental trajectories during childhood and adolescence. Neuroimage 36, 1065–1073.

Molfese, P.J., Glen, D., Mesite, L., Cox, R.W., Hoeft, F., Frost, S.J., Mencl, W.E., Pugh, K.R., Bandettini, P.A., 2021. The Haskins pediatric atlas: a magnetic-resonance-imaging-based pediatric template and atlas. Pediatr Radiol 51, 628–639.

Morris, S.R., Holmes, R.D., Dvorak, A.V., Liu, H., Yoo, Y., Vavasour, I.M., Mazabel, S., Madler, B., Kolind, S.H., Li, D.K.B., Siegel, L., Beaulieu, C., MacKay, A.L., Laule, C., 2020. Brain Myelin Water Fraction and Diffusion Tensor Imaging Atlases for 9-10 Year-Old Children. J Neuroimaging 30, 150–160.

Najdenovska, E., Aleman-Gomez, Y., Battistella, G., Descoteaux, M., Hagmann, P., Jacquemont, S., Maeder, P., Thiran, J.P., Fornari, E., Bach Cuadra, M., 2018. In-vivo probabilistic atlas of human thalamic nuclei based on diffusion-weighted magnetic resonance imaging. Sci Data 5, 180270.

Namdari, A., Li, Z., 2019. A review of entropy measures for uncertainty quantification of stochastic processes. Advances in Mechanical Engineering 11.

Palmer, C.E., Pecheva, D., Iversen, J.R., Hagler, D.J., Jr., Sugrue, L., Nedelec, P., Fan, C.C., Thompson, W.K., Jernigan, T.L., Dale, A.M., 2021. Microstructural development from 9 to 14 years: Evidence from the ABCD Study. Dev Cogn Neurosci 53, 101044.

Pauli, W.M., Nili, A.N., Tyszka, J.M., 2018. A high-resolution probabilistic in vivo atlas of human subcortical brain nuclei. Sci Data 5, 180063.

Rapuano, K.M., Laurent, J.S., Hagler, D.J., Jr., Hatton, S.N., Thompson, W.K., Jernigan, T.L., Dale, A.M., Casey, B.J., Watts, R., 2020. Nucleus accumbens cytoarchitecture predicts weight gain in children. Proc Natl Acad Sci U S A 117, 26977–26984.

Sanchez, C.E., Richards, J.E., Almli, C.R., 2012. Age-specific MRI templates for pediatric neuroimaging. Dev Neuropsychol 37, 379–399.

Van Hecke, W., Leemans, A., Sage, C.A., Emsell, L., Veraart, J., Sijbers, J., Sunaert, S., Parizel, P.M., 2011. The effect of template selection on diffusion tensor voxel-based analysis results. Neuroimage 55, 566–573.

Wells, W.M., 3rd, Viola, P., Atsumi, H., Nakajima, S., Kikinis, R., 1996. Multi-modal volume registration by maximization of mutual information. Med Image Anal 1, 35–51.

White, N.S., Leergaard, T.B., D’Arceuil, H., Bjaalie, J.G., Dale, A.M., 2013a. Probing tissue microstructure with restriction spectrum imaging: Histological and theoretical validation. Hum Brain Mapp 34, 327–346.

White, N.S., McDonald, C., Farid, N., Kuperman, J., Karow, D., Schenker-Ahmed, N.M., Bartsch, H., Rakow-Penner, R., Holland, D., Shabaik, A., Bjornerud, A., Hope, T., Hattangadi-Gluth, J., Liss, M., Parsons, J.K., Chen, C.C., Raman, S., Margolis, D., Reiter, R.E., Marks, L., Kesari, S., Mundt, A.J., Kane, C.J., Carter, B.S., Bradley, W.G., Dale, A.M., 2014. Diffusion-weighted imaging in cancer: physical foundations and applications of restriction spectrum imaging. Cancer Res 74, 4638–4652.

White, N.S., McDonald, C.R., Farid, N., Kuperman, J.M., Kesari, S., Dale, A.M., 2013b. Improved conspicuity and delineation of high-grade primary and metastatic brain tumors using “restriction spectrum imaging”: quantitative comparison with high B-value DWI and ADC. AJNR Am J Neuroradiol 34, 958-964, S951.

Wilke, M., Holland, S.K., Altaye, M., Gaser, C., 2008. Template-O-Matic: a toolbox for creating customized pediatric templates. Neuroimage 41, 903–913.

Wu, G., Jia, H., Wang, Q., Shen, D., 2011. SharpMean: groupwise registration guided by sharp mean image and tree-based registration. Neuroimage 56, 1968–1981.

Xie, W., Richards, J.E., Lei, D., Zhu, H., Lee, K., Gong, Q., 2015. The construction of MRI brain/head templates for Chinese children from 7 to 16 years of age. Dev Cogn Neurosci 15, 94–105.

Yoon, U., Fonov, V.S., Perusse, D., Evans, A.C., Brain Development Cooperative, G., 2009. The effect of template choice on morphometric analysis of pediatric brain data. Neuroimage 45, 769–777.

Zhang, S., Arfanakis, K., 2013. Role of standardized and study-specific human brain diffusion tensor templates in inter-subject spatial normalization. J Magn Reson Imaging 37, 372–381.

Zhao, T., Liao, X., Fonov, V.S., Wang, Q., Men, W., Wang, Y., Qin, S., Tan, S., Gao, J.H., Evans, A., Tao, S., Dong, Q., He, Y., 2019. Unbiased age-specific structural brain atlases for Chinese pediatric population. Neuroimage 189, 55–70.

Zhu, J., Zhang, H., Chong, Y.S., Shek, L.P., Gluckman, P.D., Meaney, M.J., Fortier, M.V., Qiu, A., 2021. Integrated structural and functional atlases of Asian children from infancy to childhood. Neuroimage 245, 118716.

Zhuang, J., Hrabe, J., Kangarlu, A., Xu, D., Bansal, R., Branch, C.A., Peterson, B.S., 2006. Correction of eddy-current distortions in diffusion tensor images using the known directions and strengths of diffusion gradients. J Magn Reson Imaging 24, 1188–1193.

Fan, C.C., Palmer, C.E., Iversen, J.R., Pecheva, D., Holland, D., Frei, O., Thompson, W.K., Hagler, D.J., Andreassen, O.A., Jernigan, T.L., Nichols, T.E., Dale, A.M., 2022. FEMA: Fast and efficient mixed-effects algorithm for population-scale whole-brain imaging data. bioRxiv, 2021.2010.2027.466202.

