## Supplemental Information for "Multimodal Image Normalisation Tool (MINT) for the Adolescent Brain and Cognitive Development study: the MINT ABCD Atlas"

Supplementary Table 1. Summary of previously published pediatric atlases.

| Publication | No. of subjects | Age (years) | Sex (% female) | Race/ethinicity | Reg modality | Atlas images |
| --- | --- | --- | --- | --- | --- | --- |
| Molfese et al., 2021 | 72 | 7-14 (mean =10) | 44% | Caucasian (*n*=66),  African American (*n*=4),  Asian (*n*=1)  Hispanic (*n*=1)  mixed-race (*n*=1) | T1 (acquired on 1.5T) | T1, Freesurfer ROIs. |
| Wilke 2008 | 404 | 4-18 | 52% | Not given | T1 (acquired at 1.5T) | T1 |
| Xie et al 2015 | 138 | 7-16 | 37% | Chinese | T1 (3T) | T1 |
| Zhao et al 2019 | 328 | 6–12 | 47% | Chinese | T1 (3T) | T1, T2, tissue probability maps |
| Avants 2015 | 119 | 7-18 | 51% | Matched the demographic distribution of children ages of 7 to 18 years in the United States, based on US Census data, in terms of race, ethnicity, gender and family income. | T1 | T1, DTI, arterial spin labelling perfusion magnetic MRI, blood oxygen |
| Morris et al 2020 | 20 | 9-10 | 20% | Not given | T1 | T1, DTI, myelin water imaging |
| Fonov et al 2011 | 324 | 4-18 | ? | Representative of the U.S. population with respect to income (as a proxy for socioeconomic status) and race/ethnicity, | T1 | T1, T2, and proton density-weighted images |
| Yoon et al 2009 | 53 | 8-9 | 57% | Not given | T1(1.5T) | T1, T2, and proton density-weighted images |
| Sanchez et al 2012 | 1289 in total, varies accordiging to age group (median group size 31) | 4-24, divided in to 32 age groups at 6 month increments | 50% overall, varies by age group (median for age groups 51%) | Not given for full cohort or separate age groups | T1(1.5T) | T1, T2 |
| Zhu et al 2021 | 272 | 7.26 – 7.92 | 52% | Chinese 48.9%  Malay 31.3%  Indian 16.9% | T1, DTI | T1, DTI, fMRI |

Supplementary Figure 1. Associations between RNI and BMI thresholded at Bonferroni corrected p-value=0.05.

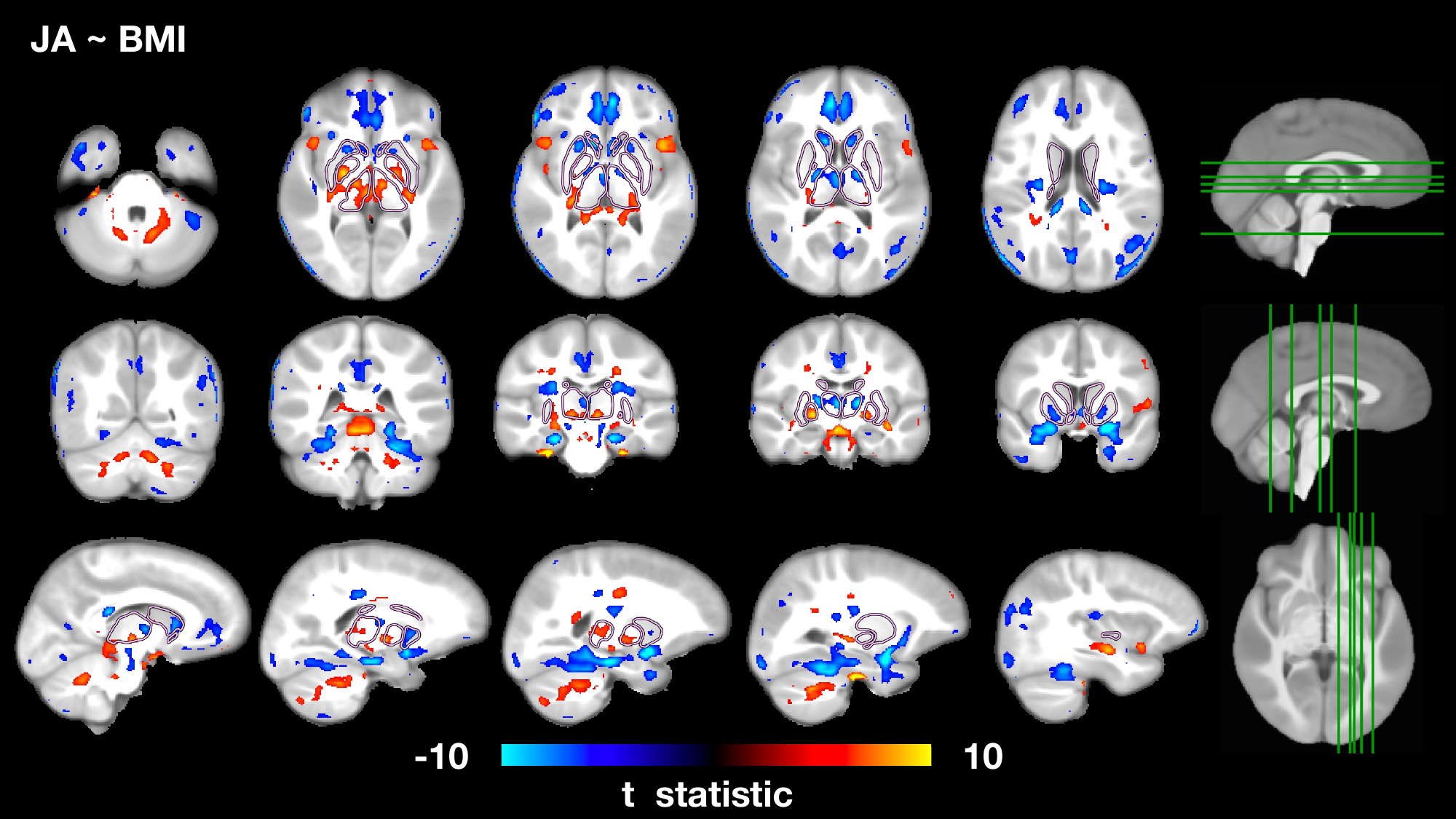

Supplementary Figure 2. Associations between JA and BMI thresholded at Bonferroni corrected p-value=0.05.
